# Muscle-specific Keap1 deletion enhances force production but does not prevent inactivity-induced muscle atrophy in mice

**DOI:** 10.1101/2024.10.03.616570

**Authors:** Edwin R. Miranda, Justin L. Shahtout, Shinya Watanabe, Norah Milam, Takuya Karasawa, Subhasmita Rout, Donald L. Atkinson, William L. Holland, Micah J. Drummond, Katsuhiko Funai

## Abstract

Immobilization-associated muscle atrophy and weakness appear to be driven in part by oxidative stress. Nuclear Factor Erythroid 2-Related Factor 2 (NRF2) is a critical redox rheostat that regulates oxidative stress responses, and its deletion is known to accelerate muscle atrophy and weakness during aging (sarcopenia) or denervation. Conversely, pharmacologic activation of NRF2 extends mouse lifespan and attenuates sarcopenia. Similarly, deletion of Kelch-like ECH-associated Protein 1 (Keap1), negative regulator of NRF2, enhances exercise capacity. The purpose of this study was to determine whether muscle-specific Keap1 deletion is sufficient to prevent muscle atrophy and weakness in mice following 7-days of hindlimb unloading (HU). To test this hypothesis, control (Ctrl) and tamoxifen inducible, muscle-specific Keap1 knockout (mKO) mice were subjected to either normal housing (Sham) or HU for 7 days. Activation of NRF2 in muscle was confirmed by increased mRNA of NRF2 targets thioredoxin 1 (Txn1) and NAD(P)H quinone dehydrogenase 1 (NQO1) in mKO mice. Keap1 deletion had an effect to increase force-generating capacity at baseline. However, muscle masses, cross sectional area, and *ex vivo* force were not different between mKO and Ctrl HU mice. In addition, muscle 4-hydroxynonenal-modified proteins and protein carbonyls were unaffected by Keap1 deletion. These data suggest NRF2 activation improves muscle force production during ambulatory conditions but is not sufficient prevent muscle atrophy or weakness following 7-days of HU.

**Graphical Abstract:** 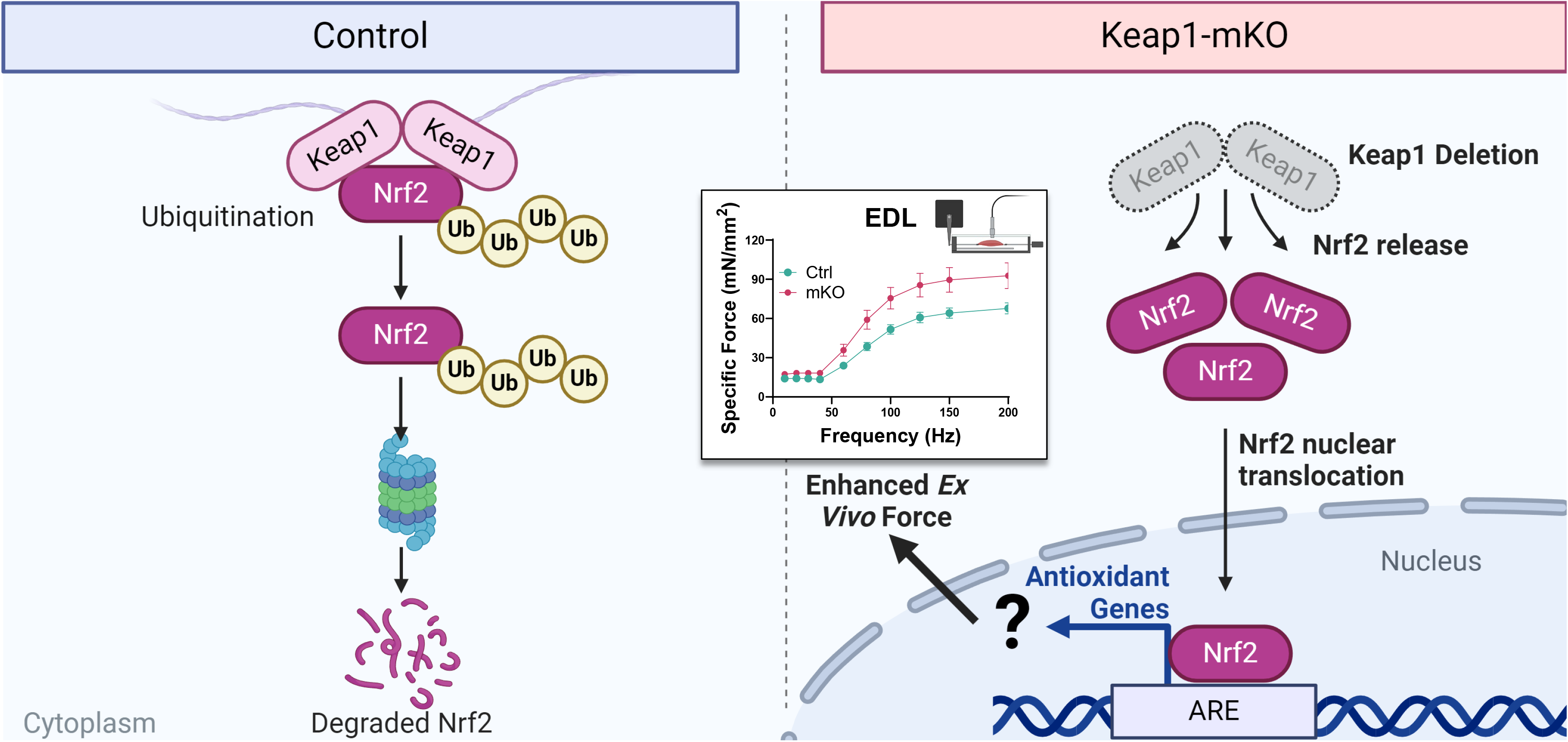

## Introduction

Maintenance of muscle mass and strength is critical for resiliency during cancer, aging, and immobility (*1*). Retention of muscle mass and function is also critical for the preservation of ability to perform activities of daily living during these processes (*2*). For example, loss of muscle mass and strength in cancer-related cachexia predicts mortality (*3*). Aging slows the rate of recovery from immobility and complete recovery is often unattainable with advanced age leading to progressive loss of muscle mass and function (sarcopenia) (*4*). Despite the broad relevance and importance for preserving muscle mass and function, a pharmacologic therapy has yet to be developed to mitigate their losses in these contexts.

Accumulation of reactive oxygen (ROS) and carbonyl species (RCS) are purported as major mechanisms driving the loss of muscle mass and strength (*5-10*). Specifically, our lab and others have recently implicated the accumulation of reactive lipid carbonyls such as 4-hydroxynonenal (4HNE) to drive muscle atrophy and weakness following inactivity models such as hindlimb unloading (HU) (*11*) and denervation (*7-9*). Muscle atrophy and weakness are mitigated by preventing the accumulation of 4HNE-modified proteins with interventions such as overexpression of glutathione peroxidase 4 (GPX4) (*11, 12*), inhibition of lysophosphatidylcholine acyltransferase 3 (LPCAT3) (*13*), and treatment with the RCS scavenger N-acetyl-carnosine (*11*) or liproxstatin (*9*).

Nuclear Factor Erythroid 2-Related Factor 2 (NRF2) is a transcription factor that is known to drive intracellular antioxidant stress response. NRF2 does so by regulating the transcription of genes involved in glutathione synthesis (e.g. glutathione reductase), antioxidant enzymes (e.g. thioredoxin-1), hypoxia response gentes (e.g. hemeoxygenase-1), and xenobiotic genes (e.g. NQO1) by binding to Antioxidant Response Element (ARE) promoter regions. Global deletion of NRF2 exacerbates age-related losses in muscle mass, strength, and accelerated frailty (*14-18*).

Conversely, treating animals with NRF2 activators such as sulphoraphane have been demonstrated to attenuate muscle dysfunction, and expand lifespan in male mice (*19, 20*). NRF2 activators such as sulphoraphane are electrophiles which act on NRF2’s negative regulator Kelch-like ECH-associated Protein 1 (Keap1).

Keap1 is an E3 ubiquitin ligase that binds to and ubiquitinates NRF2 to promote its degradation upon reduction of key cysteine residues. Electrophiles such as sulphoraphane oxidize these key residues to liberate and stabilize NRF2. Because NRF2 is extensively regulated by post-translational mechanisms, overexpression of NRF2 is likely not an effective strategy to enhance NRF2 activity. NRF2 activators are known to have off-target effects, making targeting Keap1 a potentially attractive strategy to enhance NRF2 activity. Recently, muscle-specific inducible Keap1 deletion was shown to activate NRF2 and enhance muscle performance (*21, 22*).

However, it is unknown if muscle specific deletion of Keap1 is sufficient to prevent the loss of muscle mass and strength following a bout of inactivity such as hindlimb unloading. The purpose of this study was to test the hypothesis that muscle-specific deletion of Keap1 would prevent disuse-induced oxidative stress, thereby attenuating the loss of muscle mass and strength.

## Methods

### Animal model

To study the effects of Keap1 deletion in skeletal muscle, we generated a tamoxifen-inducible skeletal muscle-specific knockout (Keap1-mKO, or mKO) mouse. Keap1-mKO mice were generated by crossing conditional Keap1 knockout mice (Keap1^lox/lox^) mice acquired from Jackson Laboratory (RRID:IMSR_JAX:037075)(*23*) with HSA-MerCreMer mice (Figure 1A). Genotypes were determined via PCR using ear punch material as the source of DNA (Figure 1B). For experiments, 8-12 week old control (Keap1^lox/lox^ without Cre) or Keap1-mKO (Keap1^lox/lox^ with Cre) mice were injected with tamoxifen (7.5 µg/g body mass) for five consecutive days (Figure 1C). After a 2-week washout period from the last day of injection, mice either underwent 7-days of hindlimb unloading (HU) or normal housing (Sham) as previously described (*11*) (Figure 1C). Mice were housed with a 12-hour light/dark cycle in a temperature-controlled room. Mice were fasted for 1 hour prior to anaesthetization via intraperitoneal injection of 80 mg/kg ketamine and 10 mg/kg xylazine and tissue harvest. All experimental procedures were approved by the University of Utah Institutional Animal Care and Use Committee.

**Figure 1.**
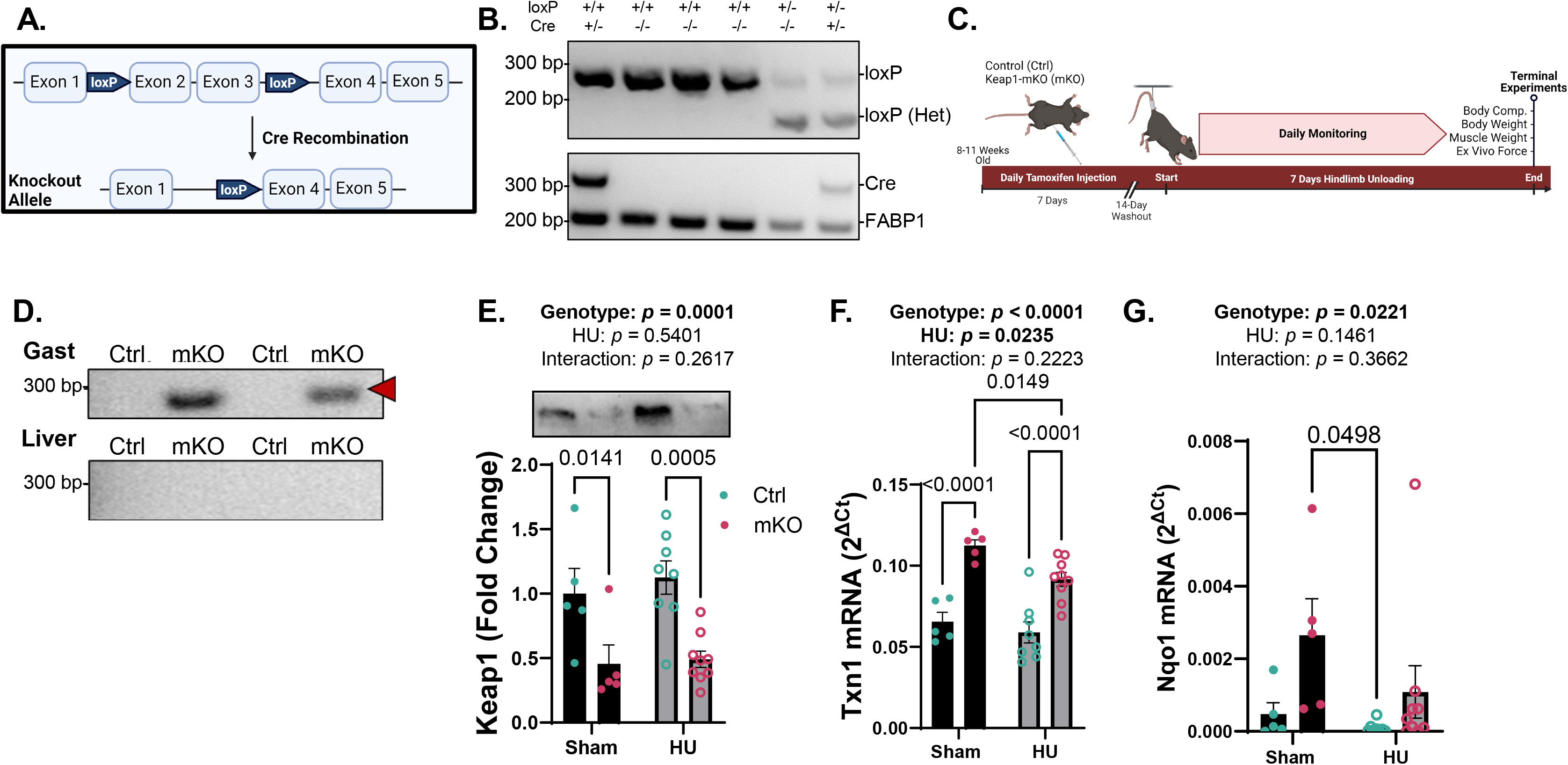
Validation of skeletal muscle specific knockout of Keap1. A. Schematic of Keap1 inducible knockout allele. B. Genotyping for loxP (top gel) and Cre (bottom gel) performed on ear snips of mice. C. Timeline of the experimental design. Ctrl and mKO male mice were injected with tamoxifen for 5 days followed by a 2-week washout before undergoing either 7-days of hindlimb unloading (HU) or normal housing (Sham) before terminal measures were assessed. D. Genotyping for successful Cre recombination following tamoxifen injection. Only successfully recombined alleles are amplified to produce a 288 bp fragment (Arrow) in gastrocnemius muscle from mKO mice (top) but not in liver (bottom) or in loxP Ctrl mice. E. Western blot of Keap1 in mouse gastrocnemius from Ctrl and mKO mice. F-G. RT-PCR for NRF2 target genes Txn1 and Nqo1 in gastrocnemius muscles from Ctrl and mKO mice. Data are Mean ± SEM and analyzed via Two-way ANOVA with Bonferroni Post hoc tests. Statistical significance was set to *p*<0.05. Schematics in panels A and C were made by BioRender.

### Hindlimb unloading

A timeline of the experimental design is presented in Figure 1C. Mice underwent 7-days of HU with two mice per cage as previously described (*6, 11, 13*). These mice were checked daily to ensure food and water consumption and to measure body mass. After the 7-days of HU, body mass and composition of the mice was assessed via Nuclear Magnetic Resonance (NMR, Bruker Minispec LF50) and mice were fasted for 1 hour prior to anaesthetization and tissue harvest. The soleus (SOL), extensor digitorum longus (EDL), gastrocnemius (GAS), tibialis anterior (TA), and plantaris (PLANT) were dissected from mice and quickly weighed before either being assessed for ex vivo force production, flash frozen in liquid nitrogen, or preserved in optimal cutting temperature (OCT) compound.

### Ex vivo skeletal muscle force production

Force generating capacity of both the SOL and EDL were measured as previously described (*6, 11, 13, 24*). Briefly, SOL and EDL were tied with suture at proximal and distal tendons then attached to an anchor and force transducer in a tissue bath (Aurora Scientific, Model 801C) while being submerged in oxygenated Krebs-Henseleit Buffer solution at 37^°^ C. The optimal length of the muscle was determined via maximum twitch force production. The buffer was then replaced with fresh buffer solution and equilibrated for 5 minutes. After equilibration, a force-frequency sequence stimulated the muscle at increasing frequencies (10, 20, 30, 40, 60, 80, 100, 125, 150, and 200 Hz) with a 2-minute rest interval between each frequency. The rates of contraction and relaxation were quantified as previously described (*6, 11, 13*). The Aurora Scientific DMAv5.321 software was utilized for analysis of force production data.

### Skeletal muscle histology

OCT embedded EDL samples were mounted on a precooled platform and sectioned at 10 µm thickness with a cryostat (Microtome Plus). 6-8 serial sections from each control of mKO sample were transferred onto the same slide. Sections then underwent blocking for 1 hour at room temperature with M.O.M. mouse IgG blocking reagent (Vector Laboratories, MKB-2213), after which they were incubated overnight at 4^°^ C with a concentrated primary antibody targeting myosin heavy chain (MHC) IIa (SC.71, IgG1, 1:100, DSHB) in 2.5% horse serum in PBS. The next day, sections were incubated for 1 hour at room temperature with anti-mouse IgG conjugated to Alexa Fluor 488 (Invitrogen, A28175) (staining MHC IIa). Sections were then washed three times in PBS, fixed with methanol for 5 minutes at room temperature and then washed 2 more times in PBS. Finally, slides were mounted with mounting medium (Vector Laboratories, H-1000). Slides were imaged on a Zeiss Axioscan.Z1 at 20X magnification. Masks for fibers were generated via Cellpose (V3.0) (*25*) and then imported into ImageJ software where CSA was quantified and fluorescent intensity at 488 nm was used to assign fiber type where 488 nm positive cells were considered MHC IIa fibers and fluorescence negative cells were considered MHC IIx or IIb fibers. Type I fibers consist a negligible proportion in EDL muscles (*26*).

### Western blotting

Whole muscle lysates were utilized for western blotting. Approximately 20 mg of frozen GAS muscle was cut, weighed, and homogenized in a ground-glass homogenization tube using a mechanical ground glass pestle grinder in 20 volumes of ice-cold RIPA buffer (Thermo Scientific, 89901) supplemented with HALT protease phosphatase inhibitor cocktail with EDTA (Thermo Scientific, 78429). Homogenates were centrifuged for 15 minutes at 12,000 x g at 4^°^ C prior to protein estimation of the supernatant via BCA (Thermo Scientific, 23227). Equal protein was then mixed with Laemmli sample buffer with BME to a final protein concentration of 2 mg/mL and denatured by incubating at 95^°^ C for 5 minutes. 20 µg of protein was then loaded onto a 4-20% gradient TGX gel (Bio-Rad) and separated via electrophoresis (200 V, 30 minutes). Proteins were transferred to nitrocellulose membranes (110 V, 1 hour) on ice and then ponceau stain was placed on the membrane to visualize the membrane bound protein. Membranes were then blocked in 5% BSA in TBST for 1 hour at room temperature with rotation. Membranes were then incubated in primary antibodies targeting 4HNE (1:1000, Abcam, ab48506), p62 (1:1000), or LC3 (1:1000) in 5% BSA in TBST overnight at 4^°^ C with gentle rocking. Membranes were then washed with TBST, incubated in species-appropriate secondaries (1:5,000) in 3% milk in TBST for 1 hour at room temperature with rotation. After washing the membranes three times with TBST and once with TBS, Membranes were incubated in Western Lightning Plus-ECL (PerkinElmer) and were then imaged for chemiluminescence (BioRad). Image Lab software was used for densiometric analysis of bands resolved at the predicted molecular weights and this signal was made relative to the intensity of the ponceau stain on the membrane.

### Quantitative reverse transcription polymerase chain reaction

RNA was extracted from approximately 50 mg of TA muscles via column-based purification kit (Zymo-Research, R2050) according to manufacturer protocol. Following quantification, 400 ng of RNA was reverse transcribed using an iScript cDNA Synthesis kit (Bio-Rad). Reverse transcription PCR (RT-PCR) was performed with the Viia 7 Real-Time PCR System (Life-Technologies) using SYBR Green reagent (Life-Technologies). Data were normalized to the ribosomal protein L32 gene expression levels and then normalized to the mean of the control sham group. Primer sequences used in this study are as follows: Nqo1 forward: AGAGAGTGCTCGTAGCAGGA, Nqo1 reverse: CAGGATGCCACTCTGAATCG, Txn1 forward: GCCCTTCTTCCATTCCCTCTG, Txn1 reverse: AGGTCGGCATGCATTTGACT, Atf4 forward: CCTGAACAGCGAAGTGTTGG, Atf4 reverse: TGGAGAACCCATGAGGTTTCAA, L32 forward: TTCCTGGTCCACAATGTCAA, L32 reverse: GGCTTTTCGGTTCTTAGAGGA.

### Protein carbonyl assay

Protein carbonyls were assed via colorimetric assay per the manufacturer’s instructions (Cayman Chemical, 10005020). Briefly, 200 µg of GAST protein in 200 µL of RIPA supplemented with protease phosphatase inhibitor cocktail was incubated with 800 µL of dinitrophenylhydrazine (DNPH) in 2.5 M HCl at room temp for 1 hour with constant shaking. DNPH derivatized protein was then precipitated in trichloro acetic acid and then washed with 1:1 ethanol:ethyl acetate to remove un-bound DNPH. Pellets were resuspended in Guanidine HCl before being measured for absorbance at 370 nm. Background absorbances derived from the same samples that were treated with 2.5 M HCl without DNPH were subtracted from derivatized samples. Carbonyl content in nmol/mL was calculated by the following equation: Protein Carbonyl (nmol/mL) = [(CA)/(*0.011 µM^-1^)](500 µL/200 µL).

An aliquot of the assayed sample was then diluted 1:5 in water and then total protein was measured via BCA. Protein carbonyl content in nmol/mg tissue was then calculated by dividing the protein carbonyl content in nmol/mL by the protein concentration in mg/mL.

### Statistics

The data in figures are represented as the means ± standard error of the mean (SEM). Analyses were performed using GraphPad Prism 9.1.1 software. Figures were made with GraphPad Prism and BioRender. Statistical comparisons were made with two-way Analysis of Variance (ANOVA) with Bonferroni as post hoc analysis for multiple comparisons where appropriate.

Means were statistically significantly different if *p* < 0.05.

## Results

### Keap1 deletion successfully activates NRF2

Tamoxifen treatment successfully resulted in Keap1 loxP recombination in muscle but not in liver (Figure 1D). Likewise, skeletal muscle Keap1 protein levels were reduced in Keap1-mKO mice compared to control (*p* = 0.0001, Figure 1E). There was no effect of HU on Keap1 protein abundance in either genotype (Figure 1E). NRF2 target genes thioredoxin 1 (Txn1) (*p* < 0.0001, Figure 1F) and NAD(P)H quinone dehydrogenase 1 (Nqo1) (*p* = 0.0221, Figure 1G) were both increased Keap1 mKO mice compared to control. HU significantly attenuated the increase in Txn1 transcripts in mKO mice (*p* = 0.0235, Figure 1F) and trended to do the same to Nqo1 transcripts (*p* = 0.0221, Figure 1G).

*Effects of muscle-specific Keap1 knockout on body mass, body composition, and muscle mass* As expected, mice that underwent 7-days of HU had significantly lower body mass compared to the sham mice (*p* < 0.0001, Figure 2A). Keap1-mKO mice weighed significantly lower compared to control mice in sham condition (*p* = 0.0197, Figure 2A), though this effect disappeared after HU. The differences in body mass between genotypes were entirely explained by differences in lean mass (Figure 2B). HU reduced muscle masses for almost all hindlimb muscles (*p* < 0.0001, Figure 2C-G) but genotype had no significant effect on muscle mass with or without HU.

**Figure 2.**
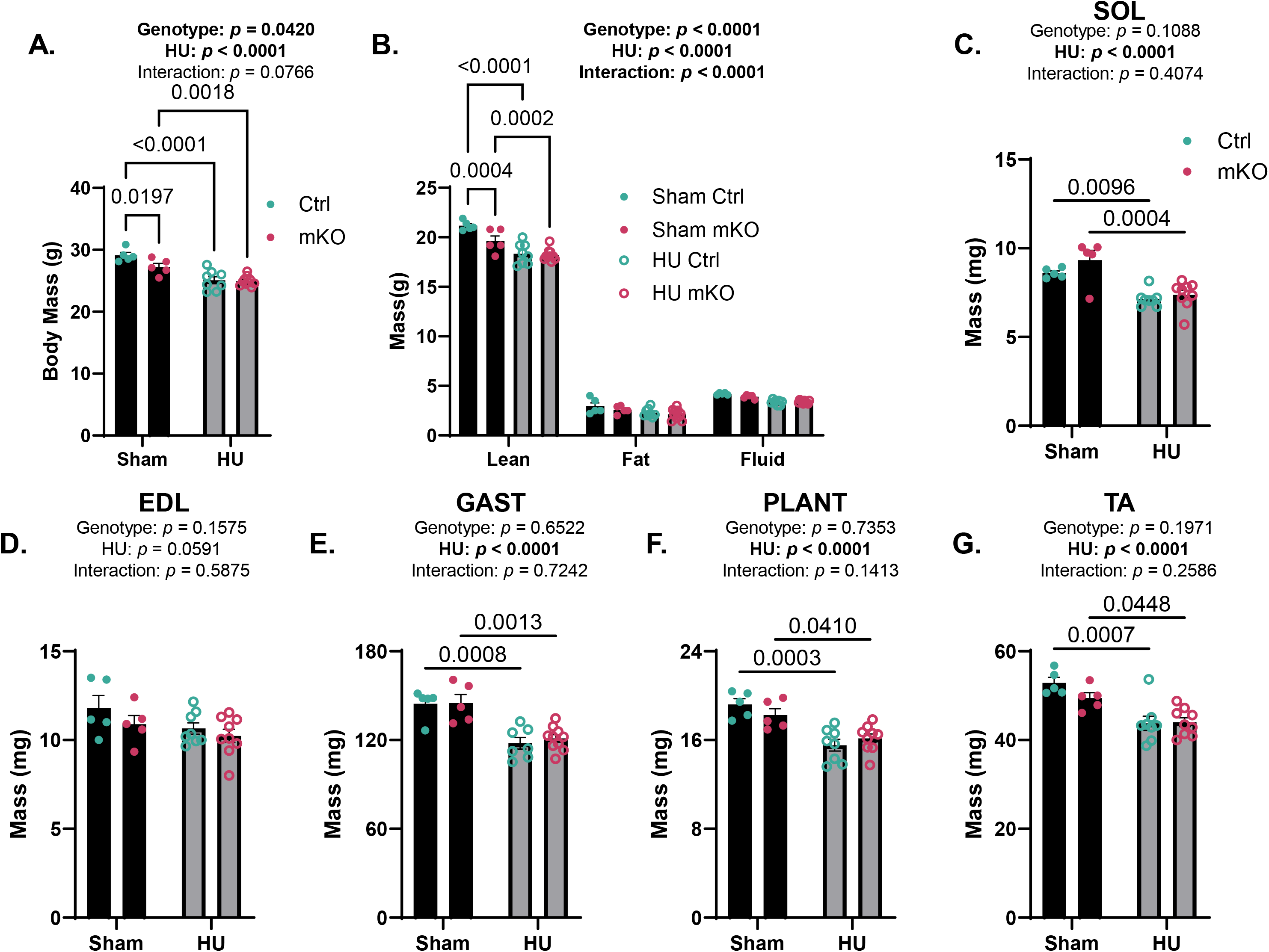
Skeletal muscle specific knockout of Keap1 does not prevent muscle atrophy following 7-days of hindlimb unloading. A. Post-intervention body mass was lower following HU and in Sham mKO mice B. Body composition via NMR reveals lower lean mass in Sham mKO mice and in both Ctrl and mKO mice following HU. C-G. Average of left and right hindlimb muscle masses taken from mice at time of dissection were lower following HU but not altered by mKO. Data are Mean ± SEM and analyzed via Two-way ANOVA with Bonferroni Post hoc tests. Statistical significance was set to *p*<0.05.

While genotype did not significantly influence muscle masses, Keap1 deletion may have more subtle effects on myofibers. Muscles from control and Keap1-mKO mice with or without HU were analyzed for fiber-type and cross-sectional area (CSA) (Figure 3A). Indeed, Keap1 deletion increased the frequency of fibers with greater CSA for both IIA and IIX/IIB fibers during the sham condition (Figure 3B and 3C), though this effect disappeared after HU. Additionally, there was a trend for a fiber-type switch from IIA to IIX/IIB with Keap1 deletion only with HU.

**Figure 3.**
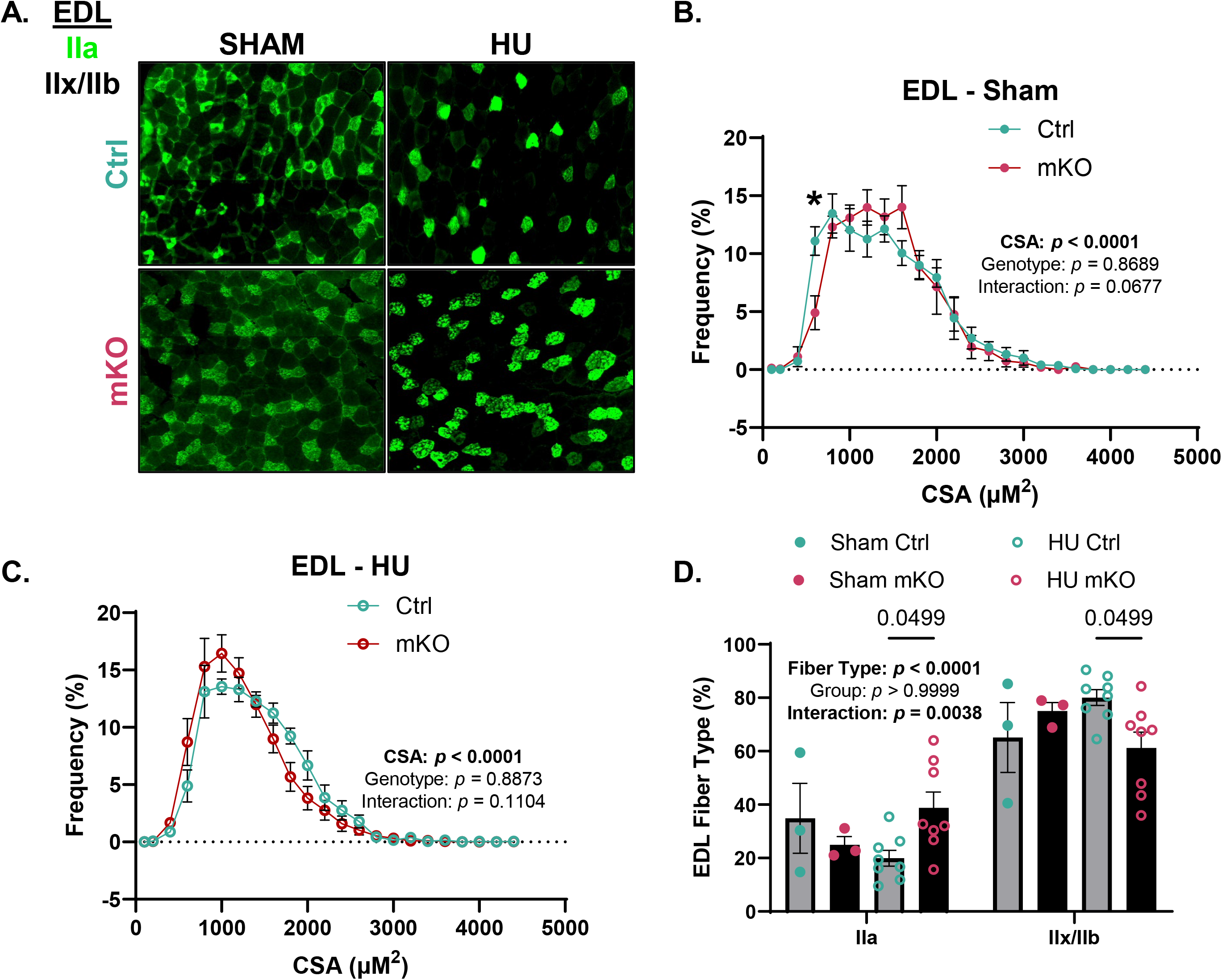
Skeletal muscle specific knockout of Keap1 decreases fast twitch fiber CSA following HU. A. Representative image of EDL cross section and immunofluorescence targeting Type IIa fibers (green). B. Frequency of EDL fiber cross sectional area in Sham and HU (C) were not different between genotypes. D. Fast, oxidative IIa fibers represent a greater proportion of EDL muscles in HU mKO mice compared to HU Ctrl mice. Data are Mean ± SEM and analyzed via Two-way ANOVA with Bonferroni Post Hoc tests. Statistical significance was set to *p*<0.05. * Indicates Post hoc significance.

*Keap1 deletion improves skeletal muscle force-generating capacity in ambulatory mice* Hindlimb unloading is known to induce reduction in force-generating capacity in addition to loss of skeletal muscle mass (*6, 11*). We quantified the force production of soleus and EDL muscles *ex vivo* and analyzed them as both absolute force (mN) and normalized to muscle area (specific force, mN/mm^2^). In sham condition, Keap1 deletion significantly improved absolute and specific force in EDL (Figure 4A and 4B), with but not in soleus (Figure 4C and 4D). However, the effect of Keap1 deletion on muscle force disappeared in mice that underwent HU.

**Figure 4.**
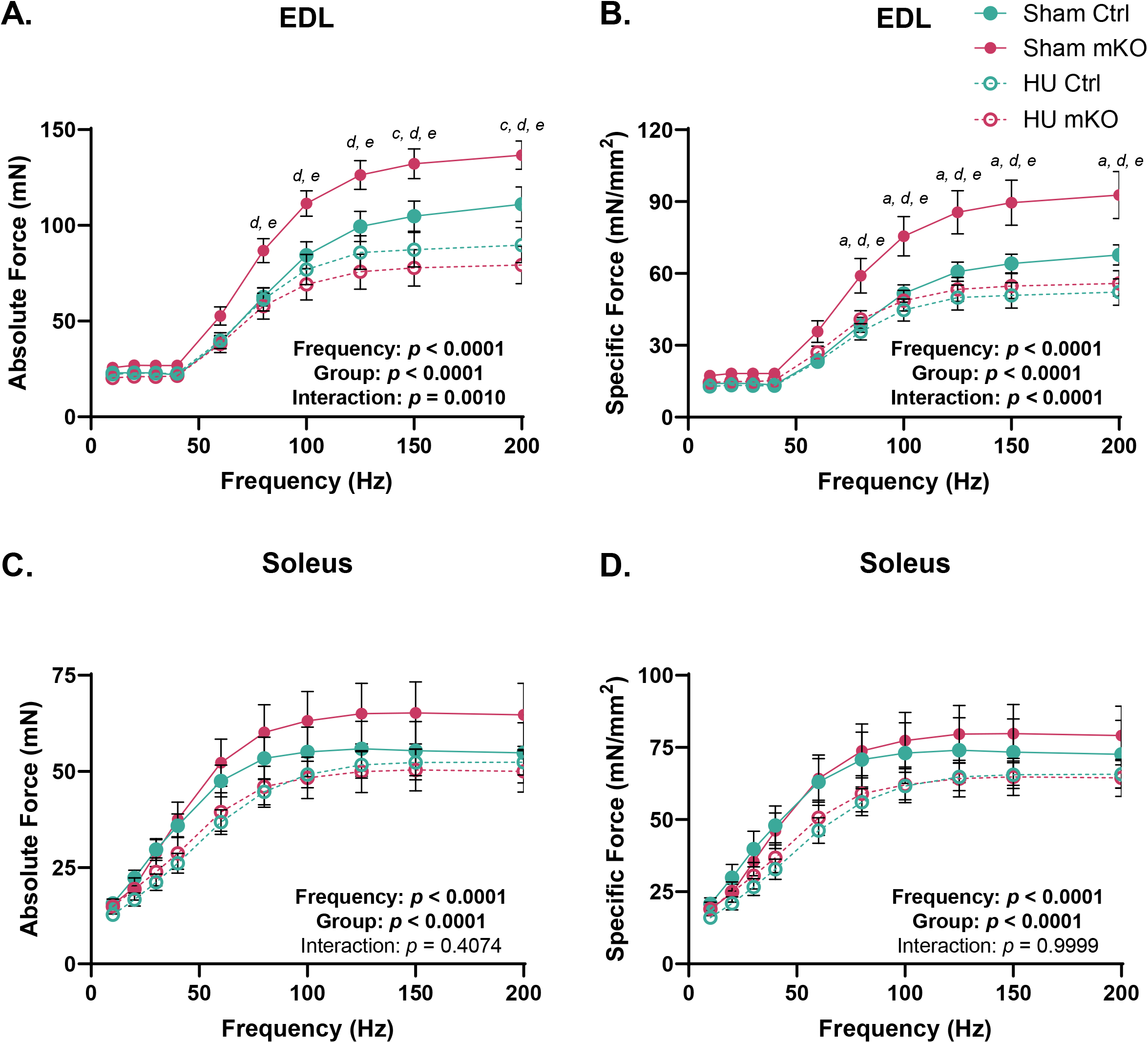
Skeletal muscle specific knockout of Keap1 enhances *ex vivo* force but does not prevent muscle weakness following 7-days of hindlimb unloading. A. Force frequency curves quantifying absolute *ex vivo* force in soleus muscles was higher in Sham mKO mice but was no different between Ctrl and mKO mice following HU. Specific force was calculated by normalizing *ex vivo* force production to muscle cross sectional area. B. Soleus *ex vivo* specific force was higher in Sham mice but was not affected by Keap1 knock out. *Ex vivo* absolute (C) and specific force (D) were highest in mKO sham mice. Data are Mean ± SEM and analyzed via Two-way ANOVA with Bonferroni Post hoc tests. Statistical significance was set to *p*<0.05. Post Hoc significance is indicated as follows: *a* – Sham Ctrl v Sham mKO, *b* – Sham Ctrl v HU Ctrl, *c* – Sham Ctrl v HU mKO, *d* – Sham mKO v HU Ctrl, *e* – Sham mKO v HU mKO, *f* – HU Ctrl v HU mKO.

### Effect of muscle-specific Keap1 knockout on stress response signaling

To assess muscle carbonyl stress, 4HNE-modified proteins were assessed in gastrocnemius muscles via western blot and total protein carbonylation was assessed via colorimetric assay. Contrary to our hypothesis, neither HU nor Keap1 influenced levels of 4HNE-modified proteins (Figure 5A and 5B) or global protein carbonylation (Figure 5C). Given that Keap1 is known to interact with p62, a known regulator of autophagy, we also assessed the abundance of LC3I, LC3II, and p62 in muscles from control and Keap1-mKO mice with or without HU. Both LC3I (*p* = 0.0034, Figure 5D) and LCII (*p* =0.0084, Figure 5E) were elevated in response to HU with no effect of Keap1 deletion. LC3II/I ratio, was not significantly altered by HU indicating proportional increases in LC3I and LC3II in response to HU (Figure 5F). Protein abundance of p62 was also unaltered by HU or by Keap1 deletion (Figure 5G). Lastly, we assessed the mRNA of Atf4, an alternative stress response regulator. Similar to other stress indicators we measured, Atf4 mRNA was also not perturbed by HU or Keap1 deletion in muscle (Figure 5H).

**Figure 5.**
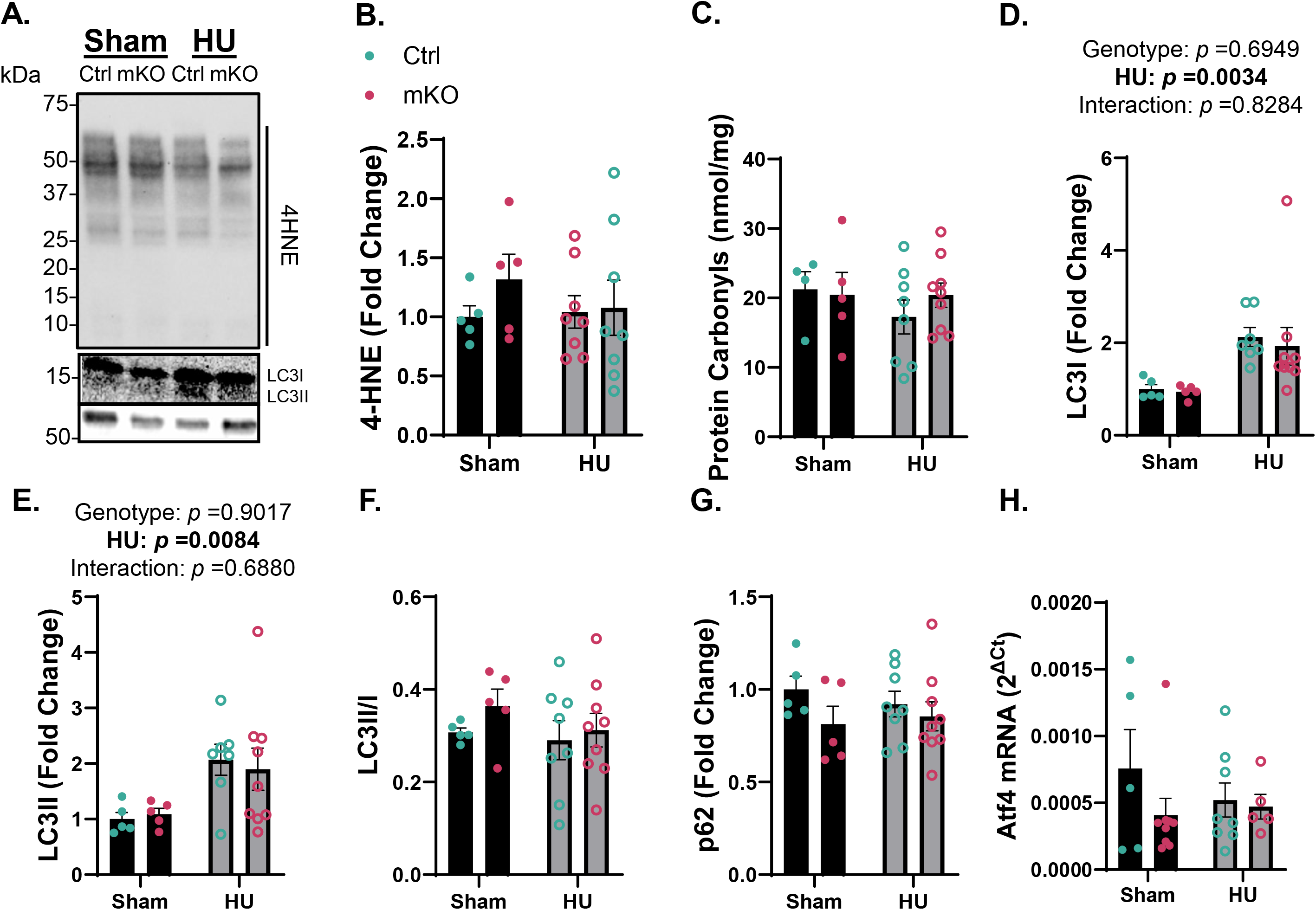
Skeletal muscle specific knockout of Keap1 does not affect autophagic flux or carbonyl stress. A. Representative western blot for LC3I, LC3II, p62, and 4HNE. B. Quantification of 4HNE western blots revealed no effect of mKO or 7-days of HU on 4HNE-modified proteins. C. Global protein carbonylation was also not altered by mKO or 7-days of HU. Both LC3I (D) and LC3II (E) protein abundance was not affected by mKO but was increased following 7-days of HU. However, the proportion of LC3II:LC3I (F) was unaffected by mKO nor 7-days of HU. G. p62 was also unaffected by mKO and 7-days of HU. H. The expression of stress response gene Atf4 was unaltered by mKO or HU. Data are Mean ± SEM and analyzed via Two-way ANOVA with Bonferroni Post hoc tests. Statistical significance was set to *p*<0.05.

## Discussion

Maintenance of muscle mass and function during insults that promote their losses is predictive of recovery from injury, disease, and propensity for developing frailty during aging (*2, 4*). Lack of the antioxidant transcription factor NRF2 promotes muscle atrophy and weakness similar to that induced by denervation (*27*), and accelerates sarcopenia during aging (*14-17*). Conversely NRF2 activators and muscle specific knockout of the NRF2 negative regulator Keap1, enhances exercise capacity (*22*) and extend lifespan in mice (*20*). However, this is the first study to demonstrate that muscle-specific Keap1 deletion is insufficient to prevent muscle atrophy and weakness following 7-days of HU despite its effect on enhancing force-generating capacity in sham mice.

Muscle-specific deletion of Keap1 significantly increased *ex vivo* force in EDLs of sham mice. Keap1 deletion had no effect on fiber-type composition, but CSA distribution was slightly shifted toward smaller fibers in the control mice compared to the mKO mice. Nonetheless, this slight shift in CSA size distribution is unlikely to explain the robust differences in force production between control and mKO mice in sham condition. In contrast, there were no differences in the force-generating capacity between control and mKO mice that underwent 7-day HU intervention. This suggests, that while Keap1 deletion has the ability to improve muscle force production at baseline, it is insufficient to protect mice from HU-induced muscle weakness.

In contrast, Keap1 deletion had no influence on muscle mass with or without HU. This was despite the previous findings that NRF2 may regulate skeletal muscle mass. Keap1 knockout appeared to successfully upregulate NRF2 activity, evidenced by increased expression of NRF2 target genes Nqo1 and Txn1. Contrary to previous work, including our own (*11*), neither 4HNE nor total protein carbonyls were elevated with HU. Our lab has previously shown that 7-days of HU is sufficient to induce accumulation of 4HNE-modified proteins (*11*). 4HNE represents only a subset of reactive carbonyls, and 4HNE antibodies likely do not exhibit equal affinity to all 4HNE-conjugated proteins. Thus, it remains possible that HU elevated other RCS not recognized by this antibody. Nevertheless, it is clear that 4HNE conjugation to proteins that are detected by the 4HNE antibodies used in our study (Abcam, ab48506) is not necessary for the loss of muscle mass and function induced by HU in our mice.

Keap1 is chaperoned by p62 to the autophagosome for degradation (*28*). Therefore, we also tested whether Keap1 deletion altered p62 abundance and autophagic flux. HU resulted in a proportional increase in the abundance of both LC3I and LC3II with no effect with Keap1 deletion. There was also no effect of HU or Keap1 deletion on p62 abundance. Activation of p62 and subsequent Keap1 degradation has been shown to be an important pathway for the activation of NRF2 in response to stress (*28-30*). However, our data suggest that autophagy-mediated Keap1 degradation is not involved in skeletal muscle atrophy during HU, neither is it altered with Keap1 deletion. We also measured the stress response transcription factor Atf4 (*31*) given its ability to heterodimerize with other transcription factors such as NRF2 (*32*) and its potential role in muscle atrophy (*33*). Similar to other markers of stress we assessed, Atf4 mRNA expression was not altered by HU nor by Keap1 deletion. Collectively, these data suggest that young mice possess the intrinsic capacity to adequately respond to HU-induced stress to prevent the dramatic accumulation of carbonyl stress and dramatic activation of stress response pathways.

It is important to note that current studies were performed in young mice, and that Keap1 deletion may have protective effects in disuse in old mice that lack sufficient antioxidant capacity

(*34*). Alternatively, it is also possible that NRF2 activation in muscle cells is simply ineffective to restore muscle function with immobilization stress such as HU. Similarly, we previously demonstrated that a mouse model overexpressing mitochondria-directed catalase (a gene regulated by NRF2) is insufficient to protect them from HU-induced muscle atrophy (*6*). Paradoxically, deletion of another NRF2 target CuZn-Superoxide Dismutase (SOD1) (*5*) and NRF2 (*14, 16*) accelerate sarcopenia. Thus, it is possible that these redox mechanisms of muscle atrophy are more relevant in the context of aging rather than acute immobility. Another possibility is that Keap1/NRF in other cell types abundant in whole muscle tissue, such as endothelial cells, satellite cells, resident macrophages, and fibro-adipogenic precursors (FAPS), are important for muscle mass during immobility (*35*). Beyond NRF2, Keap1 is known to associate with other proteins such as p62 (*36*). Nonetheless, p62 protein abundance was not affected with Keap1. Finally, chronic activation of NRF2 may induce reductive stress in some contexts leading to negative outcomes (*37*). Thus, it is possible that intermittent activation of NRF2 will demonstrate more optimal protective stress response.

In conclusion, muscle-specific deletion of Keap1 enhanced EDL *ex vivo* force production but was not sufficient to prevent atrophy or weakness in mice following 7 days of HU. These findings suggest that chronic activation of NRF2 may not be effective in preventing disuse-induced reduction in skeletal muscle mass and force-generating capacity. It remains possible that targeting Keap1/NRF2 axis could be protective in other muscle wasting conditions induced by oxidative stress, particularly for age-associated decline in muscle function. It is also likely that Keap1/NRF2 axis may be play in other cell-types that reside in the skeletal muscle tissue.

Finally, future studies should also consider a possibility that intermittent activation of NRF2 might be a more appropriate approach in controlling redox state to suppress muscle atrophy and weakness.

